# Host-shift adaptation shapes genome architecture in *S. eubayanus*

**DOI:** 10.64898/2026.01.12.698964

**Authors:** Pablo Villarreal, Juan I. Eizaguirre, Carlos A. Villarroel, Tomás A. Peña, Nicolas Agier, Christian I. Oporto, Valentina Abarca, Pablo A. Quintrel, Catalina Muñoz-Tapia, Felipe Muñoz-Guzmán, Quinn K. Langdon, Chris Todd Hittinger, Javier Echeverría, Roberto F. Nespolo, Gilles Fischer, Diego Libkind, Francisco A. Cubillos

## Abstract

The host environment can profoundly shape the genome architecture of microbial species, and *Saccharomyces eubayanus*, the wild progenitor of lager yeast, provides a natural system to study this process. Most populations are associated with *Nothofagus* trees across Patagonia, whereas related Holarctic strains occur in the Northern Hemisphere and are associated with non-*Nothofagus* hosts. The evolutionary events leading to the emergence of these northern populations on a novel host remain unclear. Here, we analyzed 471 genomes from eight countries and different hosts, *Nothofagus* in the Southern Hemisphere and non-*Nothofagus* tree species in the Northern Hemisphere. Phylogenomic analysis identified eight Patagonian lineages and revealed that Holarctic strains derived from recent admixture among Patagonian ancestors, generating the genomic background of the lager-yeast mother lineage. Long-read assemblies showed that non-*Nothofagus*-associated strains harbor an elevated burden of structural variants (SVs), particularly in subtelomeric *MAL* and *IMA* regions, involved in sugar metabolism. Phenotypic tests confirmed that *Nothofagus* isolates efficiently metabolize maltose, while non-*Nothofagus* strains do not, a pattern linked to recurrent SVs and loss-of-function mutations in *MAL33*. Consistently, bark-sugar profiling revealed that maltose is abundant in *Nothofagus* but absent in non-*Nothofagus* hosts, providing an ecological context for these genomic and phenotypic differences. These results support a model in which northward dispersal of Patagonian lineages into non-*Nothofagus* forests enriched admixed genotypes, generating genomic mosaics that accumulated structural changes and losses in maltose utilization. This interplay between gene flow and genome flexibility enabled host switching and global expansion, illustrating how ecological transitions reorder genomes and drive microbial diversification.

## INTRODUCTION

Under natural conditions, physiological changes occur gradually over time, driven by the interaction of genetic variation and environmental pressures^1^, in a process by which biotic and abiotic selective forces can promote the fixation of beneficial mutations, favoring advantageous genotypes and thereby driving adaptive evolution^2,3^. In general, natural populations harbor an extensive array of genomic polymorphisms, including single-nucleotide polymorphisms (SNPs) and large-scale modifications such as chromosomal rearrangements and genome duplications, collectively shaping populations’ adaptability and survival^4–7^. Dissecting how genomic changes, including large-scale structural variants, shape populations’ capacity to cope with fluctuating environments is key to understanding the origin of biodiversity and the resilience of ecosystems^8,9^.

Large-scale resequencing projects have transformed our understanding of population genomics, revealing millions of single-nucleotide polymorphisms and uncovering the global structure, diversity, and evolutionary trajectories of natural populations^4^. Complementing these insights, advances in third-generation single-molecule long-read sequencing have significantly increased our understanding of genome structure and complexity, providing exceptional opportunities to decipher the genetic and molecular bases of adaptation in natural environments^5,8^. This technology enables the discovery of genome plasticity, the capacity of genomes to generate structural variations (SVs) such as copy number changes, insertions, deletions, and chromosomal rearrangements in response to their environment^10^. Such changes, often undetectable with short-read sequencing, can reshape genome structure, influence traits, and promote adaptation. Together, these approaches highlight both fine-scale (SNP-level) and large-scale (SV-level) dimensions of genomic variation, thereby connecting molecular mechanisms with evolutionary processes^5,11^.

Beyond point mutations, large-scale chromosomal rearrangements, aneuploidy, and horizontal gene transfer have been shown to promote adaptation in both natural and human-influenced environments^5,12–15^. These genomic alterations can influence gene dosage, disrupt or create regulatory circuits, and enable the acquisition of novel metabolic capacities. Genomic alterations are frequently shaped by natural selection, promoting lineage-specific adaptation to specific ecological niches. Indeed, host-associated habitats can exert strong selective forces on microbial populations, driving genomic divergence and metabolic specialization^16^. Such genomic flexibility, ranging from point mutations to large-scale rearrangements, enables rapid adaptation to diverse ecological settings and often drives lineage-specific divergence under strong selective pressures. This evolutionary dynamism is particularly evident in wild yeast species that occupy distinct environmental niches, including the cold-adapted *Saccharomyces eubayanus*. This species is mostly found on *Nothofagus* trees in Patagonia and on other trees such as Oak (Quercus), Pine, and Silver Maple in the Northern Hemisphere^17–19^. These hosts differ markedly in their biogeography and bark chemistry, providing contrasting ecological contexts that can shape yeast adaptation^20^. Consistent with these ecological differences, strains isolated from *Nothofagus* and non-*Nothofagus* trees display contrasting phenotypic profiles across several carbon sources, including glucose, galactose, and maltose^18^.

In *S. cerevisiae*, structural variants have been shown to explain more phenotypic diversity than SNPs across lineages^4^. However, in wild yeasts, the relative contribution of SVs to ecological differentiation remains poorly understood. Addressing this gap is essential to explain how wild yeast lineages associated with different hosts have diverged, admixed, and ultimately contributed to the emergence of lager yeast. Also, this furnishes *S. eubayanus* as a unique biological model for studying ecological adaptation, given its dual host associations and worldwide distribution. In this context, SVs and the population history of *S. eubayanus* may provide a mechanistic basis for understanding host divergence, thus explaining the yeast’s ability to adapt to the different ecological contexts, represented by *Nothofagus* and other non-*Nothofagus* species. Studying these structural genomic features, therefore, offers important insights into the mechanisms driving yeast adaptation to diverse environmental and host-associated niches.

In this study, we combined newly generated and publicly available data to assemble a dataset of 471 *S. eubayanus* genomes, providing a comprehensive view of a wild yeast. By integrating population structure, host associations, and phylogenetic analyses, we uncovered tree-associated populations, genomic signatures of adaptation, and the timing of admixture and migration events. Our results reveal that most lineages are restricted to the Patagonia region. In contrast, Holarctic strains are admixed derivatives of Patagonian populations, representing the wild ancestors that recently contributed to the origin of lager yeast. Finally, we demonstrate that structural genomic variation, together with admixture, is the key driver event of population differentiation and may underlie the ecological associations of *S. eubayanus* with *Nothofagus* and other hosts.

## METHODS

### Yeast strains

Most strains and genome sequences (n = 283) used in this study were previously reported^17,18^ and are indicated in **Table S1**. An additional 188 *S. eubayanus* strains were obtained from 23 sampling sites (**Table S1**). Briefly, bark samples from *N. pumilio, N. dombeyi,* and *Quercus rubra* trees were obtained as previously described^18^ and incubated in 10 mL enrichment media (2% yeast nitrogen base, 1% raffinose, 2% peptone, and 8% ethanol)^21^. After incubation, colonies were processed as described by Nespolo et al.^16,18^.

### Whole genome sequencing and variant-calling

yHQL strains were sequenced as previously described by Kominek, J et al.^22^. For the other strains re-sequence in this study, DNA was extracted using a Qiagen Genomic-tip 20/G kit (Qiagen, Hilden, Germany). The library prep reaction was a 100x miniaturized version of the Illumina Nextera method. Samples were sequenced on a NextSeq 500/550 High Output Kit v2.5 (300 Cycles) flow cell. Reads were processed with fastp 0.19.4 (-l 37–3)^23^. Reads were aligned against the *S. eubayanus* CBS12357^T^ reference genome^24^ using BWA-mem^25^. Duplicate reads were removed with the ‘samtools markdup’ tool^25^. Variant calling and filtering were done with GATK HaplotypeCaller version 4.1.8^26^ (–sample-ploidy 2, –min-base-quality-score 20). Functions CombineGVCFs and GenotypeGVCFs were used to merge output VCF files. We only considered SNPs with no missing data across all datasets using the *vcftools*^27^ option max-missing 1 and a maximum of 2 alleles (max-alleles 2).

### Population genomic analyses

A VCF file containing an SNP dataset of 14,004 variants was converted to *phylip* format and used as input for IQ-TREE^28^ to generate a maximum likelihood phylogeny with 1,000 ultrafast bootstrap replicates and an ascertainment bias correction (-st DNA -o CL1105.1 -m GTR+ASC -bb 1000). The number of parsimony-informative sites was 13,873. Interactive Tree of Life (http://itol.embl.de) was used for visualization and annotation. For fastSTRUCTURE analysis, we used PLINK v1.90b6.18^29^ to select SNPs from the VCF file in approximate linkage equilibrium (pairwise r^2^<0.2, within 50-SNP sliding window and shifting every 10 SNPs), yielding a subset of 7,363 variants. The VCF file was then thinned to include only sites separated by a minimum of 1,000 bp to avoid linked sites. In total, 13,958 similarly spaced SNPs, from only *S. eubayanus* strains were used for further analysis. A principal component analysis (PCA) was performed on this dataset using SmartPCA^30^. Nucleotide diversity (π) and neutrality test Tajima’s D within each population, and genetic differentiation among populations (*F_ST_*), were calculated using the R package PopGenome^31^. Genetic distances to the reference genome were estimated directly from the variant call format (VCF) file as the proportion of single-nucleotide polymorphisms (SNPs) detected per strain relative to the total number of callable genomic sites. This metric provides a direct measure of sequence divergence from the reference genome.

### Migration events inference

A tree-based method was employed to reconstruct historical connections among the studied populations and examine gene flow utilizing the TreeMix software^32^. Initially, the analysis focused on an LD-filtered subset of *S. eubayanus* individuals that showed no evidence of recent admixture (including PA, PB-1, PB-2, PB-3, PB-4, PB-5, PB-6, and NoAm), with *Saccharomyces uvarum* serving as the outgroup. TreeMix was executed ten times for each migration event value (m) ranging from 1 to 10 (-noss –k 500), and the optimal m value was estimated using the optM R package (https://cran.r-project.org/web/packages/OptM/index.html). The model’s predictive accuracy was assessed using log-likelihood. Subsequently, TreeMix was run 100 times to generate a consensus tree and bootstrap values using the BITE R package^33^. The same procedures were applied to a dataset that included both pure and admixed variants subpopulations.

### HMW-DNA extraction and long-read sequencing

Twenty-two strains representative of the different lineages were selected to be sequenced with Oxford Nanopore Technology (ONT) (**Table S2**). High molecular weight DNA was extracted using a Qiagen Genomic-tip 20/G kit (Qiagen, Hilden, Germany) as previously described by^12^. Following the manufacturer’s protocol, libraries were sequenced using a Minion on an R10.4 flow cell (Oxford Nanopore Technologies, UK). The raw pod5 files were transformed to fastq files and debarcoded using Dorado v0.7.2 with the “super high accuracy” model (https://nanoporetech.com/accuracy).

Barcodes and adapter sequences were trimmed using Porechop (https://github.com/rrwick/Porechop) and filtered with Filtlong (https://github.com/rrwick/Filtlong) using a Phred score of 15. In addition, we used publicly available paired-end Illumina sequence data previously generated for each strain^17,18^.

### Genome assembly, annotation and SV detection

Genome assembly was performed with Fly v2.9 (--nano-hq -g 400m)^34^. Additionally, three rounds of nanopolish (https://github.com/jts/nanopolish) and pilon (https://github.com/broadinstitute/pilon) were performed. The raw assembly was polished using Illumina reads filtered with a Phred score of 30 (Burrows-Wheeler Aligner). The genome assembly was annotated using the LRSDAY pipeline^35^ and the *S. eubayanus* CBS12357^T^ reference genome as the training model. To identify the SVs in *S. eubayanus* strains, we performed pairwise comparisons between the 21 strains’ *de-novo* long-read assemblies. Using MUM&Co v3.8^36^ we detected seven types of SV: deletions, insertions, duplications, contractions, inversions, and translocations. The pipeline used MUMmer v4^37^ to perform whole-genome alignments and detect SVs ≥ 50 bp. RIdeogram package was used to visualize and map the SVs in the reference genome *S. eubayanus* CBS12357^T^ ^38^.

### Population-based molecular dating

To infer divergence times among *S. eubayanus* lineages, we used an approximation based on the population-based molecular clock framework. Estimates were obtained using the mutation parameter θw, defined as θw = 2Nθμ, where Nθ corresponds to the number of generations since the most recent common ancestor of a given lineage pair, and μ is the per-base mutation rate. This formulation allows genetic diversity to be translated into temporal estimates under a neutral model of sequence evolution^39^. Since empirical estimates of mutation rates are not available for *S. eubayanus*, we adopted the single-base mutation rate reported for *S. cerevisiae* (μ = 1.84 × 10⁻¹⁰ mutations per nucleotide per generation)^40^. This value has been widely used in population genomic studies of yeasts, under the assumption that spontaneous mutation rates are broadly conserved across *Saccharomyces* species^41,42^. Estimates were calculated using genome-wide polymorphism data from the selected strains, rather than a restricted set of loci, ensuring that θw reflects average neutral diversity across the genome.

To translate generational divergence into absolute time, we assumed a range of 1–8 mitotic divisions per day^39^, consistent with empirical and theoretical estimates of yeast growth in natural environments. Importantly, this range reflects uncertainty in wild yeast life history rather than experimental growth conditions. Our reported divergence intervals, therefore, correspond to the upper bound of this generation-rate range; assuming fewer divisions per day, whether due to nutrient availability, temperature, or other environmental factors, would proportionally shift divergence times further into the past. Consequently, the estimated dates should be interpreted as conservative, order-of-magnitude approximations rather than precise chronological values. While this approach does not explicitly model demographic complexity or selection, it provides a population-genetic timescale that is comparable to previous yeast studies and is appropriate for relative comparisons among lineages.

### Phenotypic Diversity among S. eubayanus Populations

To assess the phenotypic diversity across populations, we evaluated growth and biomass production under different pH levels (2 to 6) and maltose concentrations (2, 5, 10, and 20%) using high-throughput phenotyping with 96-well microculture plates. In summary, cells were precultured in 200 μL of YPD medium at 20°C, without agitation, for 48 hours. During the experimental phase, each well was inoculated with 10 μL of pre-inoculum to achieve an optical density (OD) of 0.06–0.2 in 200 μL. The culture OD was recorded at 620 nm every 30 minutes for up to 96 hours. We estimated four parameters from this data: lag phase, growth rate (μmax), carrying capacity (ODmax), and Area under the curve (AUC) using the Growthcurver package with its default settings^43^.

Bark composition was measured from *Quercus robur*, *Q. palustris* and *Q. suber* samples obtained from the Botanical Garden at Universidad Austral de Chile (-39.80364, -73.25061). *Nothofagus pumilio* and *N. dombeyi* samples were obtained from Nevados de Chillan (-39.90484, -71.39367). For this, 3 g of representative bark samples from tree individuals where yeasts were obtained were pulverized using a Coffee Crushing Machine for 1 minute. Aqueous extracts (10 mg dry bark/ 1 mL water) as previously described^44^. Then, 20 μL of filtered aqueous extract was injected into a Shimadzu Prominence HPLC (Shimadzu, USA) equipped with a Bio-Rad HPX–87H column, using 5 mM sulfuric acid as the mobile phase at a flow rate of 0.6 mL/min, as previously described^45^, with minor modifications. The samples were injected in triplicate (n = 3).

## Statistical analysis

All statistical analyses utilized biological triplicates. A *p*-value of <0.05 was used to determine statistically significant differences. The comparison of growth rates between populations and isolation locations was conducted using a paired *t*-test. The heat map and phenotypic principal component analysis (PCA) were generated using R software^46^, specifically the "pheatmap" and “prcomp” packages from stats 3.6.0 and visualized with the "ggplot2" and “ggbiplot” packages.

## Data availability

Raw reads have been deposited in the Sequence Read Archive (SRA) database of the National Center for Biotechnology Information (NCBI) under the BioProject accession numbers listed in Table S1.

## Results

### Patagonian origins and admixture shape the global population structure of S. eubayanus

To explore the evolutionary history of *S. eubayanus*, we initially reconstructed its phylogenetic relationships and population structure from a global collection of 471 strains (**Table S1**). The collection included strains from different geographic regions, tree hosts, and related substrates: Chile (234 genomes), Argentina (206 genomes), United States (21), Canada (4), Ireland (2 genomes), China (2 genomes), New Zealand (1 genome), and two lager genomes from Denmark and the Netherlands (**Figure S1 and S2**). Illumina reads were aligned to the reference genome *S. eubayanus* CBS12357^T^, and 1,357,229 single-nucleotide polymorphisms (SNPs) were identified, corresponding to a mean SNP density of one SNP every ∼400 bp (**Table S2**). We generated a maximum likelihood phylogeny representing the *S. eubayanus* global ancestry distribution (**Figure 1A, Figure S3**).

**Figure 1.**
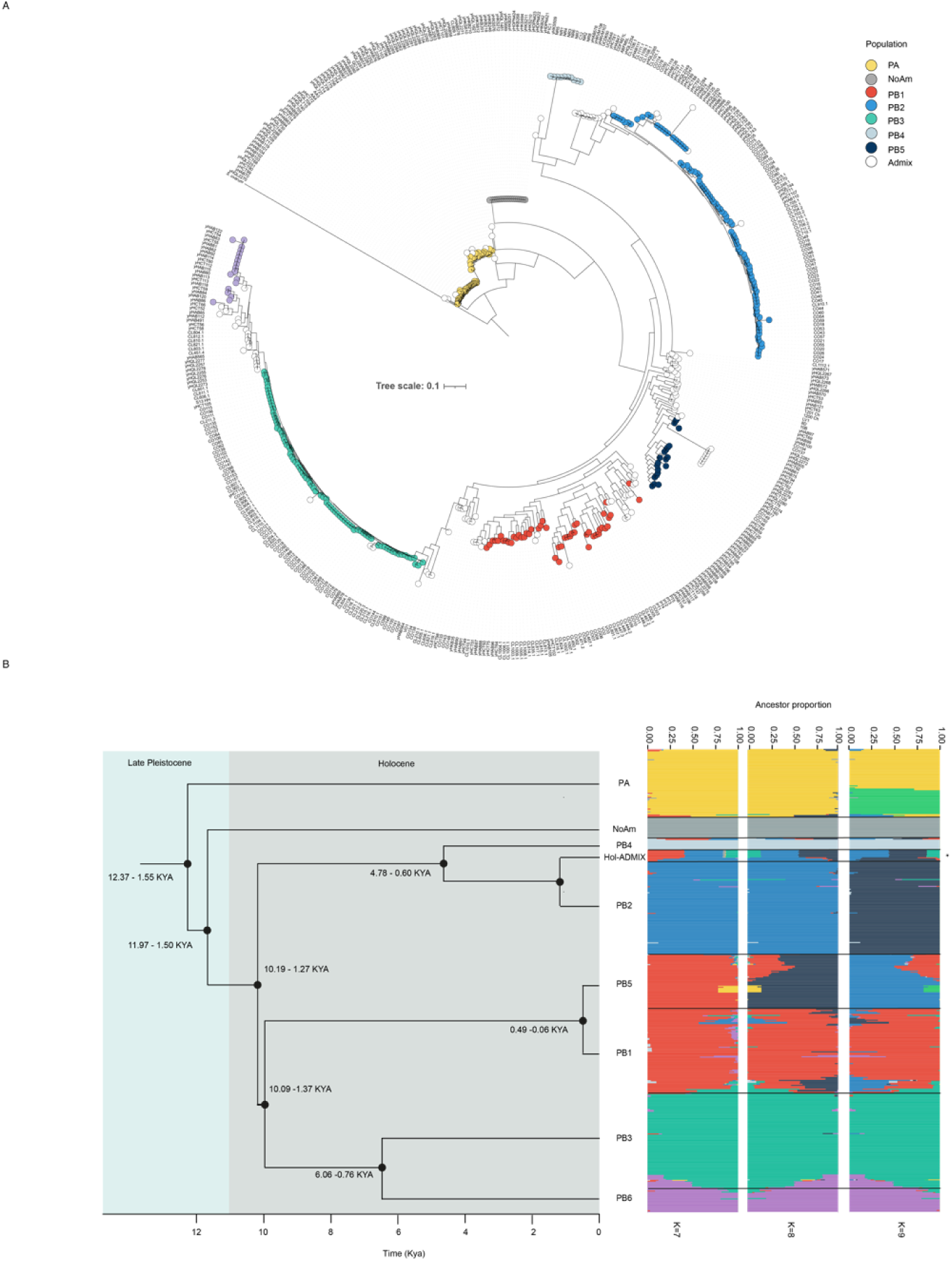
Phylogeny of *S. eubayanus*. (A) Maximum likelihood tree depicting genetic relationships between 471 strains using 14,004 biallelic SNPs (substitution model GTR+F+ASC) and manually rooted with *S. uvarum* as the outgroup. PA lineage (yellow), NoAm lineage (gray), and six PB lineages: PB-1 (red), PB-2 (blue), PB-3 (green), PB-4 (light blue), PB-5 (black), and PB-6 (purple), were identified together with admixed strains between the different lineages. White circles represent admixed strains. Branch lengths correspond to genetic distance. The tree scale is substitutions per site. (B) Estimation of the divergence time since the last common ancestor between lineages. Timeline in thousands of years (Ky). Population structure from fastStructure analysis at K = 7 to 9. Labels on the top of the bars indicate each lineage from optimum K = 8. Colored circles in (A) are representative of lineages.

In the reconstructed phylogeny, strains previously assigned to the PA lineage, mainly from Argentina, formed a well-supported sister and early-branching group (**Figure 1A**). A divergence from this node revealed North American isolates (NoAm) and a larger group of Patagonian lineages (previously referred to as PB) as distinct clusters. Within PB, strains fall into the previously identified PB-1, PB-2, and PB-3 lineages. We also identified a particular set of strains branching near PB-2, which share a recent common ancestor with this lineage. Interestingly, isolates previously classified as part of the Holarctic lineage (from North America, Asia, and Europe) are embedded within the PB-2 cluster in the current sampling, suggesting a more complex evolutionary history than previously understood. However, our sampling is skewed toward Patagonia, with limited coverage in Europe and Asia, which may limit the detection of rare Northern clades.

To determine the most likely number of lineages within the species, the genetic structure of *S. eubayanus* was determined using the fastSTRUCTURE clustering algorithm (**Figure 1B**) and Principal Component Analysis (PCA). The fastSTRUCTURE analysis revealed a clear genetic subdivision among lineages and supported an optimal number of eight genetic clusters (K = 8), as indicated by the lowest cross-validation error (Min CV: 0.279) (**Figure S4**). This analysis achieved high resolution in distinguishing the different lineages, identifying three novel clusters within the PB group, which we designated as PB-4 (n = 10), PB-5 (n = 19), and PB-6 (n = 18) (**Figure 1B**). Based on these analyses, we defined eight lineages: PA, NoAm, and PB-1 through PB-6 (**Figure 1**). Notably, all lineages except NoAm are present in Patagonia. We identified numerous admixed strains (28% of all isolates), including all samples previously assigned to the Holarctic clade. These admixed isolates share ancestry components from the Patagonian PB-2, PB-3, and PB-5 lineages (**Figure 1B**). Interestingly, individuals from the PA and PB populations were mainly collected from *Nothofagus* samples (97.5%). In contrast, isolates of the Holarctic group (n = 21) were obtained from non-*Nothofagus* hosts (secondary host), such as *Quercus* and *Pinus*, suggesting that the host tree may influence the presence of genetically distinct versus admixed lineages in natural populations (**Table S1**). The PCA further supported these results, recapitulating the main structure observed in the fastSTRUCTURE analysis (**Figure S5**). Using the total number of strains, isolates were separated according to the previously determined lineages. The individuals designated as admixed were distributed throughout the plot (**Figure S5A**). A similar analysis, excluding admixed individuals, provided a better resolution in separating the different lineages. According to this analysis, the PB-4 lineage, which is at the base of the clade where PB-2 and the Holarctic admix group emerge, is the most divergent and distinct from the rest of the PB lineages (**Figure 5SB**).

Finally, to further contextualize the global distribution of *S. eubayanus*, we estimated divergence times among lineages (**Figure 1B**). This analysis revealed that the group of strains with Holarctic ancestry emerged recently, derived from admixture events (4.78–0.60 thousand years ago, KYA) (**Figure 1B**). This pattern aligns with a recent dispersal outside of South America, likely facilitated by human dispersion. Given that Hol-ADMIX strains were primarily recovered from non-*Nothofagus* species and are closely grouped within the PB-2 ancestry, these findings support a scenario in which Northern Hemisphere admixed strains migrated from Patagonia to North America and then to Europe and beyond, or possibly via a direct route from Patagonia to Europe. Altogether, these results indicate that *S. eubayanus* is structured into eight genetic lineages, most of which are confined to Patagonia. We detected extensive admixture, particularly in isolates from non-*Nothofagus* hosts, and uncovered three new PB lineages. We found that strains from the Holarctic population represent a recent migration that has extended the species’ global distribution.

### Southern Patagonian populations harbor the greatest genetic diversity

To compare patterns of genetic variation among non-admixed *S. eubayanus* lineages and determine which harbor the greatest diversity, we assessed key population genomic metrics. We calculated genetic differentiation (*F*_ST_), nucleotide diversity (π), and Tajima’s D neutrality test for each non-admixed sub-lineage of *S. eubayanus* (**Figure 2, Table S3**). *F_ST_* pairwise comparisons revealed the lowest values between PB subpopulations, indicating lower genetic distance (**Figure 2A, Table S3A**). The PA lineage exhibited the highest *F_ST_* values among all groups, including NoAm and PB, indicating significant genetic differentiation and supporting its early divergence within the species (**Figure 2A**).

**Figure 2.**
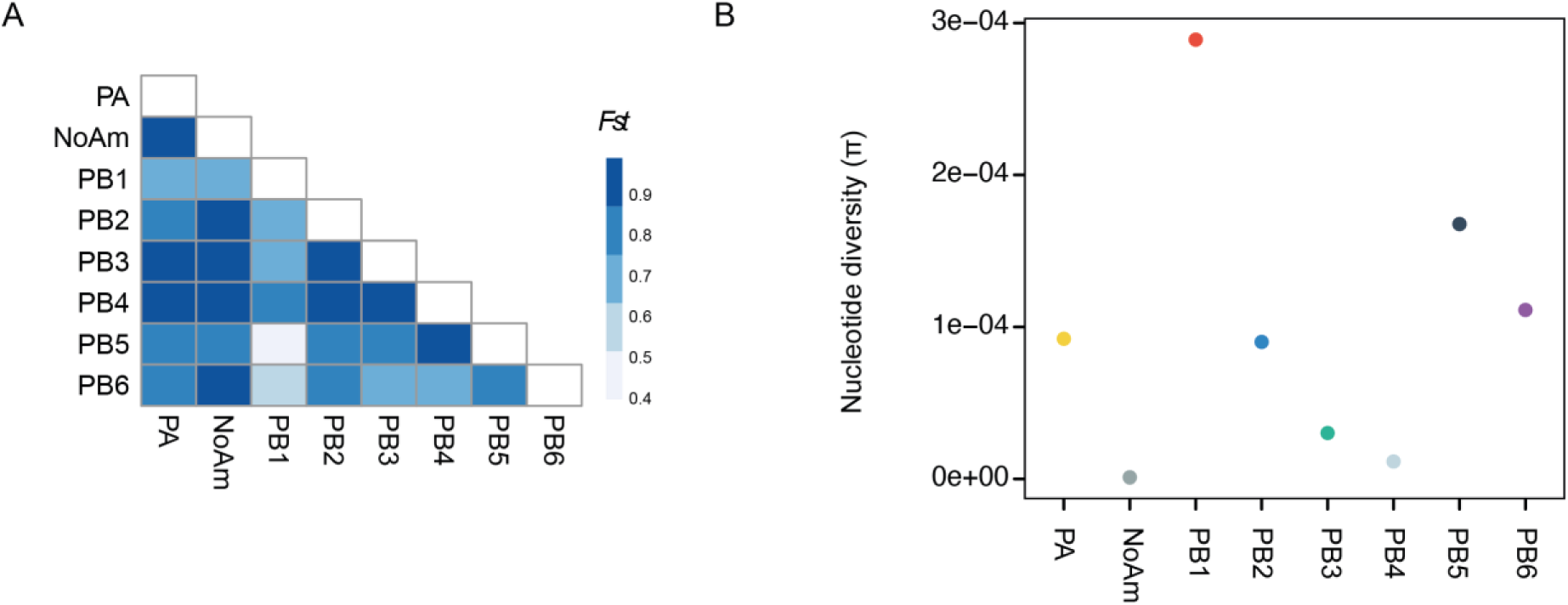
Patterns of genetic differentiation and nucleotide diversity across *S. eubayanus* lineages. (A) Pairwise *F_ST_*values between *S. eubayanus* lineages. (B) Nucleotide diversity (π) in each *S. eubayanus* lineage. Color codes in (B) represent each population identified in this work. PA lineage (yellow), NoAm lineage (gray), and six PB lineages: PB-1 (red), PB-2 (blue), PB-3 (green), PB-4 (light blue), PB-5 (black), and PB-6 (purple).

Genome-wide nucleotide diversity (π) varied nearly threefold among *S. eubayanus* lineages. The North American lineage had the lowest diversity (π = 1.06 × 10⁻⁶), while PB-1 showed the highest (π = 2.9 × 10⁻⁴). PB-6 and PB-3 also had low diversity, whereas PA, PB-2, PB-4, and PB-5 displayed intermediate levels, aligning with geographically structured diversification across Patagonia (**Figure 2B**). Tajima’s D values revealed contrasting demographic histories across Patagonia (**Figure S6, Table S3B**). Southern Patagonian lineages (PB-1, PB-4, PB-5, PB-6) showed positive D values, a pattern consistent with either balancing selection maintaining divergent haplotypes or recent population contractions reducing the frequency of rare alleles. In contrast, northern lineages (PB-2, PB-3) displayed negative values, suggesting recent demographic expansions. The PA lineage remained near neutrality, suggesting long-term stability, while the North American group showed signatures of low diversity. This interpretation aligns with an early admixture event followed by rapid expansion in novel non-*Nothofagus* environments. Overall, patterns of nucleotide diversity and Tajima’s D show that genetic variation in *S. eubayanus* is unevenly spread across lineages, reflecting the influence of geographic distribution, demographic history, and ecological factors.

### Gene flow events shape the species’ population structure

Since population structure analyses alone cannot differentiate between shared ancestry and real admixture, we investigated signals of gene flow among *S. eubayanus* populations. Our goal was to determine whether the North American (NoAm) and Holarctic (HOL) strains are truly admixed groups or separate clades, using maximum-likelihood inference in TreeMix. First, we focused on those populations that showed no admixture in our prior structure analyses, ensuring that the inferred migration edges accurately reflect genuine ancestral flows rather than recent hybrid signals. We used *S. uvarum* as an outgroup and, by modeling up to two migration events, our TreeMix prediction explained over 98% of the covariance in allele frequencies, indicating a robust fit (**Figures 3A** **and S7A**). The first edge runs from the Patagonian PA lineage into the North American clade (NoAm), and the second from PB-1 into PB-6. The presence of these edges suggests that both NoAm and PB-6 would harbor appreciable ancient gene flow (**Figure 3A**). Although we initially restricted our analysis to populations lacking admixture signals, these residual flows indicate the well-documented admixture of NoAm^17,47^ and either ancient gene exchange or unrecognized substructure within PB-6.

**Figure 3.**
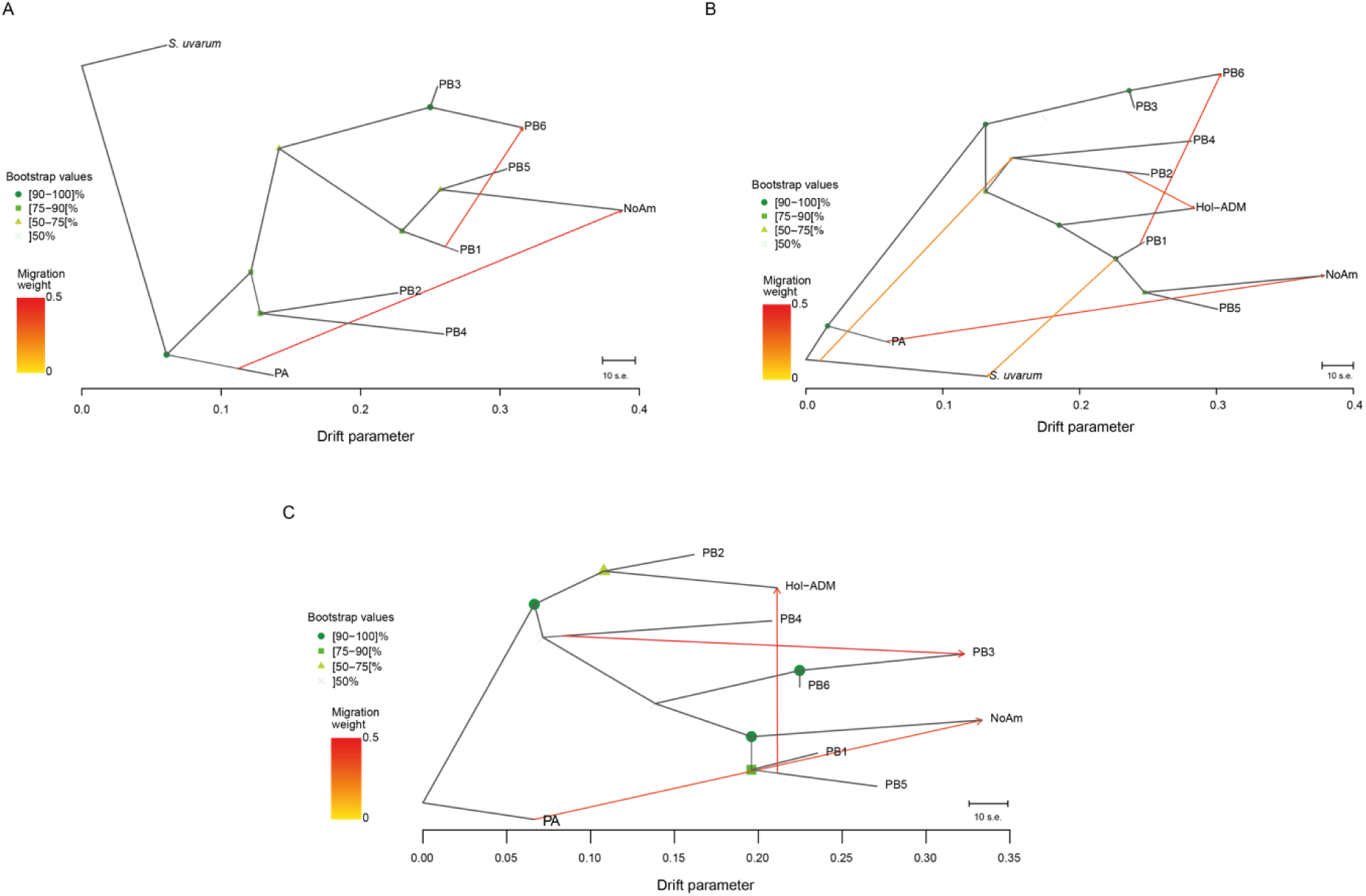
**Phylogenetic network of *S. eubayanus* lineages**. (A) Migration edges (yellow to red colored lines) estimated by TreeMix showing two migration edges on the phylogeny using pure lineages and *S. uvarum* as an outgroup. (B) Migration edges (yellow to red colored lines) estimated by TreeMix showing five migration edges on the phylogeny using pure lineages, admixed Holarctic strains, and *S. uvarum* as an outgroup. (C) Migration edges (yellow to red colored lines) estimated by TreeMix showing three migration edges on the phylogeny using pure lineages and admixed Holarctic strains. PA lineage was used as an outgroup.

Next, when we expanded the analysis to include the previously denominated Holarctic strains (**Figure 3B, Figure S7B**), three additional migration edges became apparent: first, an edge from *S. uvarum* into the ancestral node of the PB-2–PB-4 clade, suggesting ancient allelic sharing or retained polymorphism at the root of these Patagonian lineages; second, a flow from PB-1 and *S. uvarum*, which may reflect deep coalescence indicating an historical exchange; and third, a clear transfer from PB-2 into the Holarctic admixed group, hereafter referred to as Hol-ADMIX, which carries substantial Patagonian ancestry and is not a genetically distinct lineage. Finally, migration analyses using the PA lineage as an outgroup provided clearer resolution of admixture signals (**Figures 3C** and **S7C**). These analyses consistently showed gene flow from PB-5 into the Hol-ADMIX clade, in addition to the previously detected contribution from PB-2. Overall, these results demonstrate that the so-called Holarctic lineage is consistent with an admixed group with contributions from PB-2 and PB-5.

### Admixture promotes structural genome variation in Holarctic strains

To explore genomic differences between Patagonian lineages and the Hol-ADMIX group, which differ predominantly in their host trees, we selected representative strains from each lineage (PA, all PB lineages, and Hol-ADMIX), together with an admixed Patagonian strain as a control. We used ONT sequencing and assessed large-scale genomic differences between lineages. Pairwise comparisons revealed a clear pattern: Hol-ADMIX strains exhibited a higher number of structural variants (SVs) than Patagonian strains, either admixed or non-admixed (**Figure 4A**). This high number of variants shows no significant correlation with the genetic distance between strains (Spearman’s ρ = 0.43, p = 0.056, **Table S6A**). Interestingly, many Hol-ADMIX strains were isolated from a non-*Nothofagus* environment, suggesting that a secondary host may promote the accumulation or maintenance of structural variants in these genomes (**Figure 4B**). These SVs included insertions, deletions, duplications, contractions, inversions, and translocations, representing both balanced and unbalanced events, all of which showed a clear enrichment in the Hol-ADMIX strains compared to the Patagonian lineages (**Figure 4C**).

**Figure 4.**
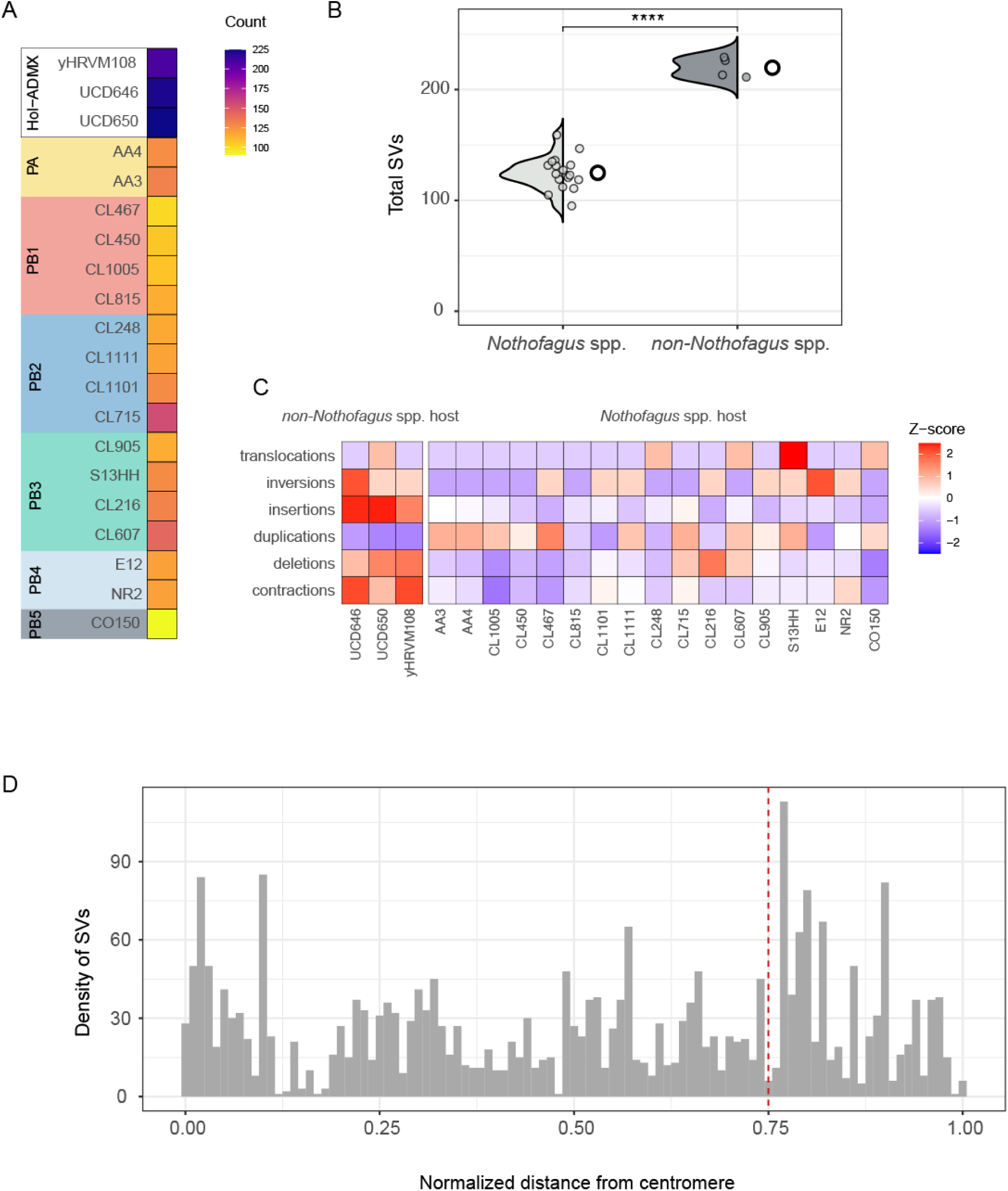
Structural variant analysis across *S. eubayanus* lineages. (A) Total number of SVs identified in each strain compared to the CBS12357^T^ reference genome. Strains are grouped and colored according to their lineage. (B) Comparison of the total number of structural variants (SVs) detected in strains isolated from *Nothofagus* spp. and non-*Nothofagus* spp. Each point represents a strain, and violin plots indicate the distribution of SV counts within each host group. (C) Z-score normalized counts of SVs, separated by SV type (insertions, deletions, duplications, contractions, inversions, and translocations). (D) Genomic distribution of unique SVs detected in the Hol-ADMIX group compared to the rest of the populations is shown as density relative to the centromere.

Mapping the SVs of Hol-ADMIX to the reference genome revealed a significant enrichment of variants in subtelomeric regions, genomic regions known for their high plasticity, dynamic gene content, and frequent rearrangements (*p*-value < 0.05, ANOVA) (**Figure 4D, Table S5 and S6B**). To further explore patterns of structural genome variation across lineages, we performed an all-vs-all comparison of the *de novo* assemblies from all sequenced strains (**Figure S8, Table S5**). This comprehensive analysis revealed that the Hol-ADMIX group consistently displayed a markedly higher number of structural variants than any of the Patagonian lineages (*p*-value < 0.05, ANOVA), even with the reference strain CBS12357, which is considered an admixture strain between Patagonian populations (**Figure S8A-B, Table S6B**). Among the different types of SVs, insertions and deletions were the most abundant (58.33% and 29.29%, respectively), accounting for most of the structural changes that distinguish admixed genomes from the Patagonian lineages (**Figure S8C**). These findings support the hypothesis that Hol-ADMIX strains harbor an excess of structural variants, particularly in subtelomeric regions, which may reflect an environmental response for genomic reorganization. Such structural variation could be a consequence of their admixed genomic background and may facilitate adaptation to a non-*Nothofagus*-associated niche.

### Maltose metabolism underlies differences between Nothofagus and *non*-Nothofagus-collected strains

To evaluate how structural variants (SVs) affect strain metabolism, we systematically examined gene families that could be disrupted by these genomic changes. Our analysis identified several families, especially the *MAL*, *IMA*, and *HXT* gene families, with SVs that likely impair their function. Interestingly, *MAL* genes were more frequently impacted in isolates from non*-Nothofagus* hosts (**Figure S9**), with recurring variants disrupting *MAL32* (maltase), *MAL31* (transporter), and *MAL33* (transcriptional activator) on Chromosome V-R (**Figure 5A, Figure S9**). A closer look at the *MAL* loci showed clear host-specific patterns: strains from *Nothofagus* maintained conserved synteny across Chromosomes V and XVI, while non*-Nothofagus*-associated isolates displayed rearrangements or complete loss of *MAL* genes, especially at the chromosome V locus (**Figure 4**). These recurring rearrangements create a consistent genomic signature that distinguishes Northern Hemisphere non*-Nothofagus*-associated admixed strains from *Nothofagus* isolates.

Phenotypic assays confirmed these differences in genomic signatures (**Figure 6**). Growth profiles across increasing maltose concentrations (2%, 5%, 10%, and 20%) showed that strains from *Nothofagus* consistently reached higher maximum optical densities and clustered according to their robust phenotype (**Figure 6, Table S7**). In contrast, strains from non*-Nothofagus* isolates displayed severely reduced growth across all tested concentrations, forming a separate phenotypic cluster (**Figure 6A, Table S7**). These results were reproducible across multiple isolates, providing consistent phenotypic separation between *Nothofagus* and non*-Nothofagus* strains.

**Figure 5.**
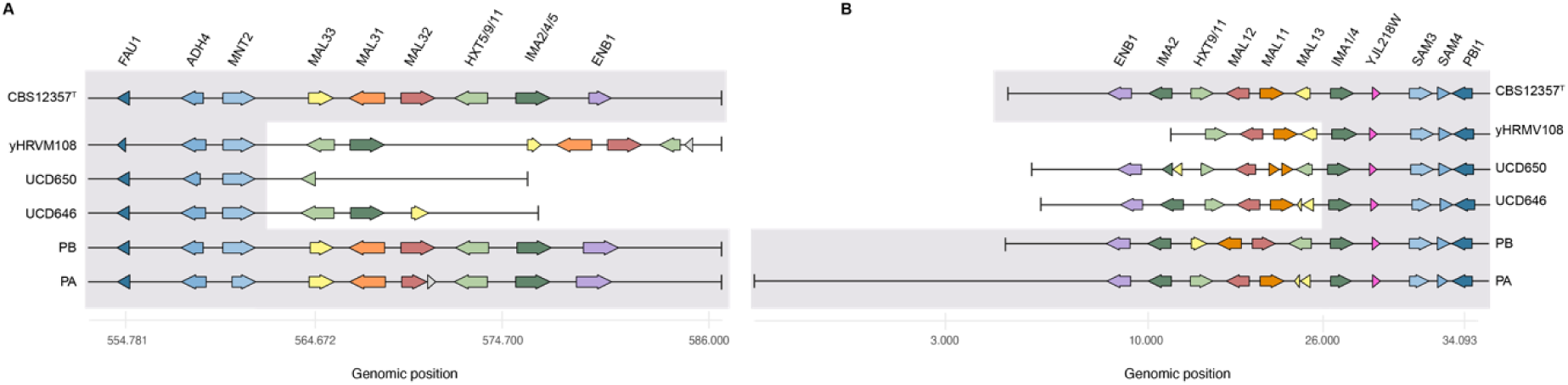
Synteny of *MAL* genes across *S. eubayanus*. (A) Genomic organization of the *MAL* locus on (A) chromosome V and (B) chromosome XVI across different lineages. Conserved synteny is shown in grey. Reference strain: CBS12357^T^. Hol-ADMIX: yHRVM108, UCD650, UCD646.

**Figure 6.**
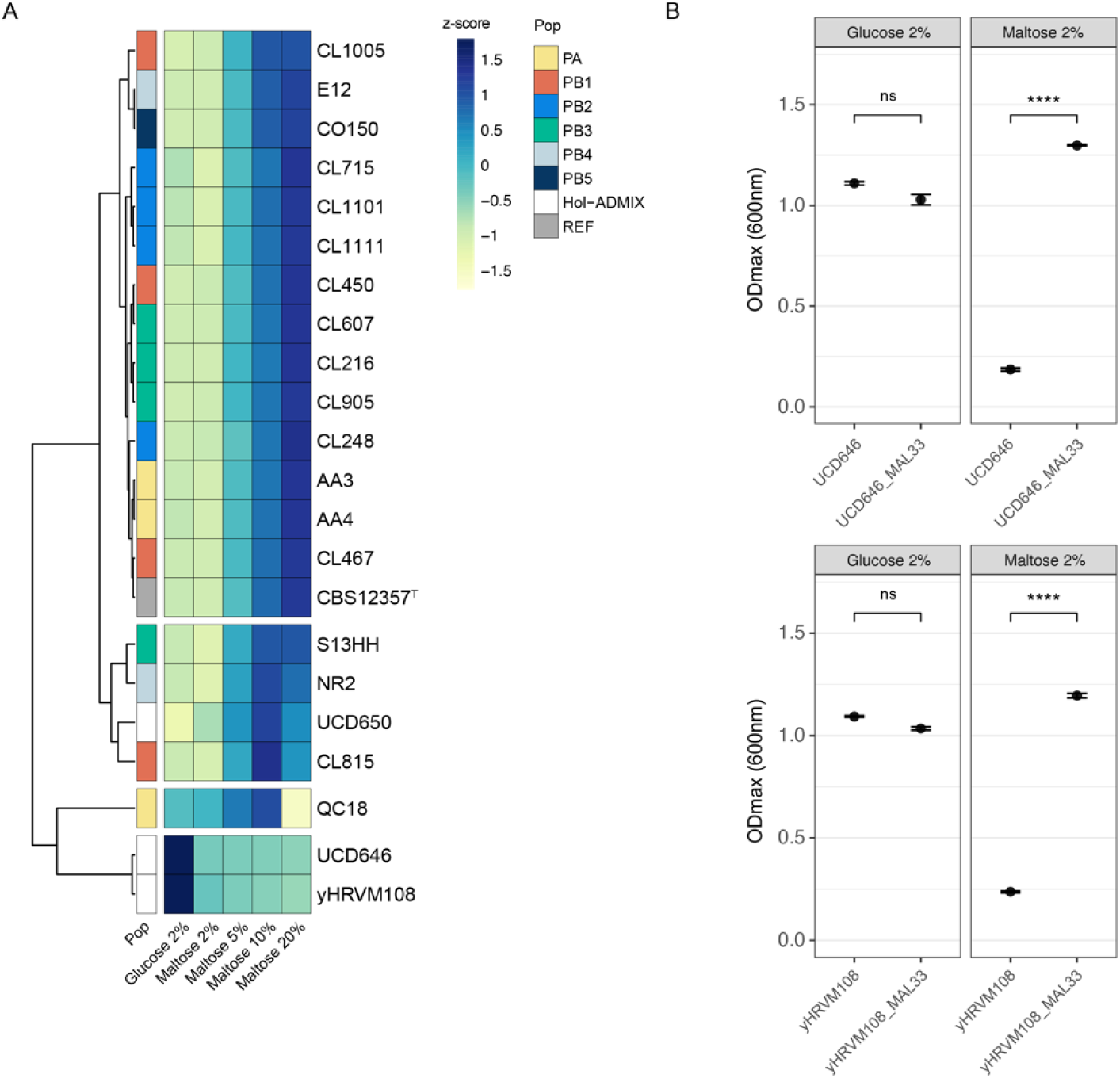
Phenotypic variation in maltose utilization among *S. eubayanus*. (A) Heatmap of z-score normalized maximum optical density of *S. eubayanus* strains across different maltose concentrations (2%, 5%, 10%, and 20%). Strains are clustered by phenotype and colored by lineages. (B) Functional validation of the MAL33 gene. Hol-ADMIX strains UCD646 and yHRVM108, which are unable to grow on maltose, were transformed with the reference CBS12357^T^ MAL33 allele, *p*-value < 0.001 (t-test). The PB CL467.1 strain was used as a control and transformed with the same plasmid.

To understand the structural differences that led to this functional divergence, we compared the organization and integrity of *MAL* loci across representative strains exhibiting contrasting maltose phenotypes (**Figures 5 and 6, Table S7**). The analysis revealed that the transcriptional activator *MAL33* was consistently affected across strains obtained from non*-Nothofagus* hosts. Strains that efficiently grew on maltose carried an intact *MAL33* gene, whereas strains unable to grow on this sugar either lacked the locus entirely or carried truncated alleles (**Figure 5**, **Figure 6A, Figure S10)**. For example, the North American isolate retained *MAL31* and *MAL32* but contained a disrupted *MAL33* gene with a premature stop codon, likely resulting in loss of regulatory function. Conversely, in the UCD strains, *MAL* genes were completely absent in chromosome V, except in UCD646, where *MAL33* was present but truncated, indicating independent structural events leading to a similar functional outcome.

To validate this hypothesis, we performed a functional complementation assay using a heterologous multicopy episomal vector (pRS426) to express the CBS12357^T^ *MAL33* allele (Promoter-ORF-terminator). This assay demonstrated that Hol-ADMIX strains isolated from non*-Nothofagus* hosts (UCD646 and yHRVM108), which were unable to grow on maltose, recovered growth when transformed with a functional *MAL33* allele (**Figure 6B, Figure S11, Table S8**). Complemented strains showed optical densities comparable to those of maltose-positive controls, confirming the contribution of *MAL33* allelic variation in the inability of non*-Nothofagus* isolates to metabolize maltose. Together, these results demonstrate that admixed non*-Nothofagus* strains lack functional maltose utilization due to recurrent structural variants in the transcriptional regulator of the *MAL* genes. In contrast, isolates obtained from *Nothofagus* trees retain a complete *MAL* loci configuration and can metabolize maltose across a range of concentrations. These contrasting genomic configurations highlight that the host environment shapes the structural architecture of *S. eubayanus* lineages, reflecting host-specific adaptive trajectories.

Finally, we examined the bark-sugar composition of *Quercus* (*Q. palustris*, *Q. suber* and *Q. robur*) and *Nothofagus* (*N. pumilio and N. dombeyi*) trees to determine whether host-related differences in carbohydrate availability could explain the contrasting genomic configurations observed among *S. eubayanus* lineages. Quantitative analyses showed clear and consistent differences in maltose concentrations between hosts. (**Table S9**). Interestingly, *Q. robur* and *Q. suber* did not show any maltose. In contrast, the different *Nothofagus* bark samples contained significantly higher maltose levels than *Q. palustris* (p-value < 0.05, ANOVA). This host-specific availability of maltose is consistent with the preservation of intact *MAL* loci in Patagonian *S. eubayanus* lineages and the recurring structural disruption of these loci in *Quercus*-associated population strains. Together, these results demonstrate a direct ecological–genomic connection between host sugar composition, structural variation at key metabolic loci, and host-shift–related adaptation in *S. eubayanus*.

## DISCUSSION

Our findings suggest that hybridization and host-dependent structural variation played a central role in the evolutionary transition of *S. eubayanus* from a Patagonian specialist to a northern-hemisphere-distributed lineage associated with habitats containing other tree species. In this alternative host environment, the wild progenitor of lager yeast experienced ecological opportunity and secondary contact among divergent Patagonian lineages, generating a hybrid genomic background that ultimately gave rise to the lager-yeast lineage. *MAL* locus remodeling in non-*Nothofagus* isolates, coupled with recurrent loss of maltose growth, supports a host-associated model of ecological differentiation. This case illustrates how environmental change can restructure genomes and alter evolutionary trajectories by creating conditions that favor the retention and sorting of hybrid variation, thereby turning hybridization from a short-lived event into a persistent evolutionary process^48,49^. In fungi^50^, plants^51^, and animals^48^, hybridization and genome admixture are well-established mechanisms that generate novel allelic combinations and structural configurations, which in turn can facilitate adaptive divergence when exposed to contrasting ecological regimes^4^.

Our genomic analyses provide additional evidence that *S. eubayanus* diversity radiates from Patagonia rather than being split between hemispheres. Earlier studies proposed an *out-of-Patagonia* dispersal model for *S. eubayanus*, with two major population groups: southern Patagonian lineages (PA and PB), and a northern Holarctic clade. By expanding the dataset to 471 genomes from both hemispheres, we uncover new Patagonian lineages and demonstrate that the Holarctic population resulted from admixture among them. Instead of forming a separate northern group, Holarctic strains descend from hybridization events among southern PB-2, PB-3, and PB-5 lineages. This admixture created a mosaic genome that later spread across the Northern Hemisphere, carrying the genetic background that ultimately led to the ancestor of lager yeast. The relatively recent timing of this event (4.78 – 0.6 KYA) suggests that human-facilitated movement created ecological pathways that allowed these admixed Patagonian genotypes to spread beyond South America^17,18^. However, given that *S. eubayanus* inhabits predominantly cold environments, where the assumptions underlying molecular clocks may not hold, this timeframe should be interpreted cautiously, and refined dating frameworks will be necessary to assess alignment with periods of human influence robustly.

A distinctive ecological pattern emerges when the natural habitats of *S. eubayanus* are examined. In *Nothofagus* forest, populations remain genetically segregated, reflecting both strong host fidelity and long-term ecological stability. In contrast, the transition into *Quercus* and other non-*Nothofagus* habitats was accompanied by hybridization among divergent Patagonian lineages, generating genomic mosaics that subsequently dispersed across North America, and Europe, and beyond^17–19^. The repeated recovery of admixed genotypes from non-*Nothofagus* bark suggests that the host plays a central role in facilitating hybridization and enabling establishment outside Patagonia. For example, Oak trees differ significantly from *Nothofagus* in their bark chemistry and sugar composition^20^ plausibly shifting the selective landscape and allowing hybrid backgrounds, rather than non-admixed lineages, to persist under these novel ecological conditions. By providing an environment in which hybrid genotypes gain a performance advantage, new northern-hemisphere tree habitats effectively transformed hybridization into a mechanism for rapid ecological accommodation, supporting the persistence and global spread of admixed populations. The Holarctic group therefore does not represent an independently evolved lineage, but instead the evolutionary outcome of host-mediated admixture and subsequent dispersal^14,15,51^.

The shift from *Nothofagus* to a secondary host in the northern hemisphere represented not just a geographical expansion but also an ecological leap that required *S. eubayanus* to adapt quickly to a chemically distinct environment. In this context, structural variants (SVs) likely provided the raw material for ecological innovation, reshaping gene content to meet new environmental challenges. SVs are well recognized as powerful engines of adaptation in fungi, as they can simultaneously affect multiple genes and regulatory regions^5,13^. Indeed, we found that SVs are particularly enriched in admixed populations associated with non-*Nothofagus* hosts, suggesting that hybridization and genome structural variation act synergistically to accelerate adaptation. Hybridization among distinct Patagonian lineages likely generated genomic mosaics containing redundant or recombined regions that became substrates for structural change. Such hybrid genomic backgrounds are known to facilitate genome plasticity and adaptive potential by unlocking hidden variation and promoting chromosomal instability^5^. Over time, beneficial rearrangements could have become fixed, conferring permanent adaptive advantages in this new ecological niche. This model would explain why SVs are disproportionately found in the admixed Holarctic group: hybridization provided a rapid genomic solution to a new host. The recurrent loss, duplication, or rearrangement of subtelomeric *MAL* and *IMA* loci in non-*Nothofagus*-associated strains exemplify the process, reflecting a functional shift away from maltose metabolism toward additional carbon sources. Comparable patterns of host-linked structural remodeling have been reported in other yeast and fungal systems, where genomic rearrangements underlie adaptation to new environments, substrates, or hosts (*S. paradoxus*, *Lachancea thermotolerans, Puccinia graminis* f. sp. *tritici, Fusarium* sp.*, and Microbotryum* sp.)^13,52–54^. Thus, structural variation emerges as a key mechanism of rapid, ecological fine-tuning, transforming hybrid genomes into adaptive prototypes capable of exploiting new and chemically challenging habitats. This evolutionary flexibility supports a dispersal model from Patagonia to the Northern Hemisphere (**Figure 7**).

**Figure 7.**
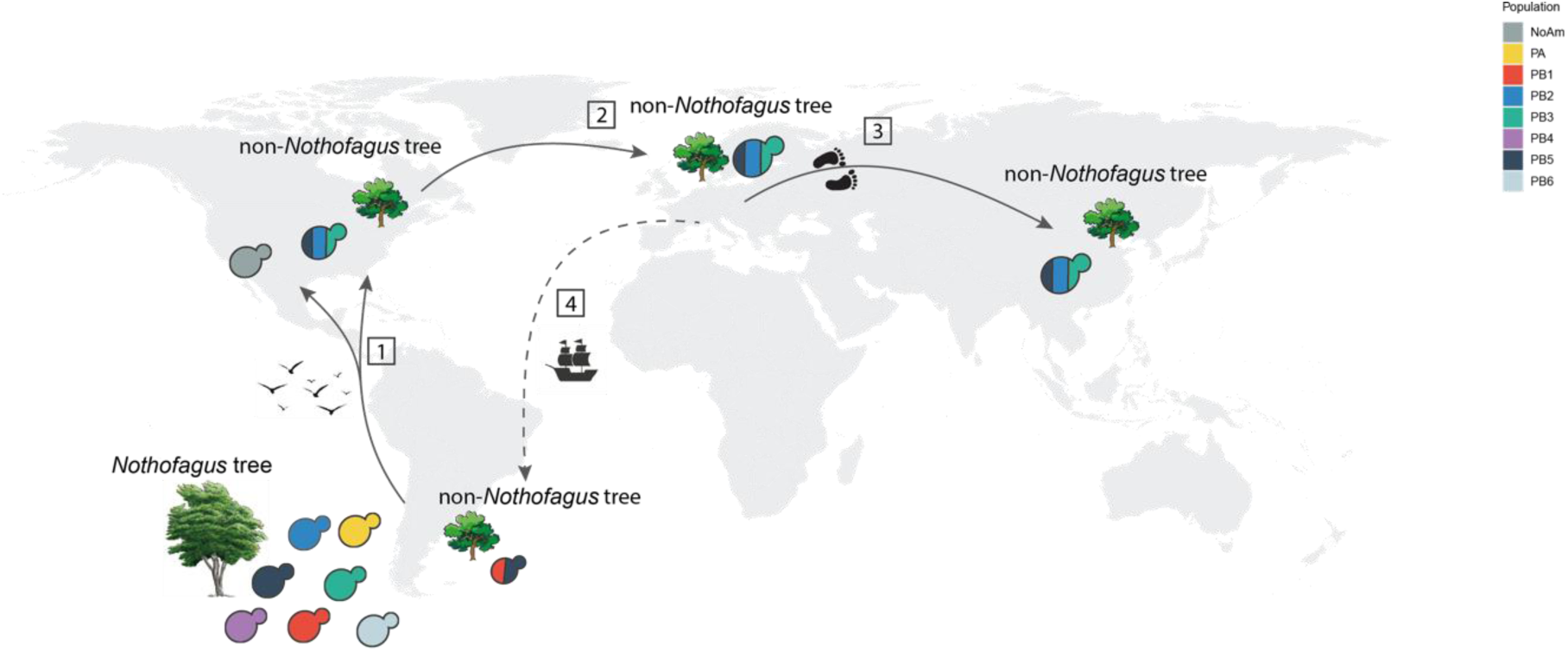
Hypothesized Model of migration and host-driven diversification of *Saccharomyces eubayanus*. (1) Multiple migrations from Patagonia to North America brought pure Patagonian lineages into non-*Nothofagus* habitats, where strong ecological filtering encouraged hybridization and genome reorganization, leading to the formation of the admixed North American (NoAm) population. (2) Later migration from North America to Europe spread mixed non-*Nothofagus*-associated strains. (3) Human-facilitated dispersal extended these lineages across Asia and other regions. (4) Introduction of Northern hemisphere tree species into South America (∼400 years ago) reconnected native Patagonian populations and introduced Holarctic lineages, sparking new hybridization events that may indicate an ongoing adaptation process within Patagonia.

Taken together, our findings reveal a dynamic evolutionary cycle in which migration, host transitions, and hybridization have repeatedly shaped the history of *S. eubayanus* (**Figure 7**). The first dispersal wave carried Patagonian lineages northward, where non-*Nothofagus* environments acted as ecological filters, selecting a subset of admixed genotypes that adapted to this novel habitat and gave rise to the North American population (1). Subsequent arrivals of additional Patagonian lineages into non-*Nothofagus* forests triggered secondary contact and hybridization, generating the admixed Holarctic strains that later spread across North America and Europe (2), including isolates recovered from China (3). Interestingly, North America harbors three different genotypes, the admixed NoAm lineage, admixed migrants with PB-1 contributions, and the Hol-ADMIX^17,55^. Human-mediated movement of *Nothofagus* or non-*Nothofagus*-associated material likely facilitated this global expansion, reinforcing the species’ *out-of-Patagonia* dispersal model. More recently, the introduction of Northern hemisphere tree species into South America has reconnected these admixed lineages with their Patagonian ancestors, marking the initiation of a new phase of local adaptation. Thus, the evolutionary trajectory of *S. eubayanus* reflects a repeating cycle of migration, admixture, and ecological innovation that began in Patagonia and continues to this day. More broadly, our findings illustrate how host transitions and genome plasticity interact to shape microbial biogeography.

## Acknowledgments

This research was funded by Agencia Nacional de Investigación y Desarrollo (ANID) FONDECYT (1220026), FONDECYT INICIACIÓN (11240649), and ANID - Programa Iniciativa Científica Milenio ICN17_022 and NCN2024_040. Research in the DL lab was supported by the National Scientific and Technical Research Council (CONICET, project PIP11220150100297), the Ministry of Science, Technology and Innovation of Argentina (MINCyT, project PICT-2020-SERIEA-00226) and the National University of Comahue (UNCo, Project 04/B247). Research in the Hittinger Lab is supported by the National Science Foundation (DEB-2110403), USDA National Institute of Food and Agriculture (Hatch Project 7005101), in part by the DOE Great Lakes Bioenergy Research Center (DOE BER Office of Science DE–SC0018409), and a Vilas Faculty Mid-Career Investigator Award.

## Author Contributions

Conceptualization: PV, DL, JIE, and FC. Methodology: PV, CAV, TAP, NA, CIO, VA, PAQ, CM and FMG. Software: PV, CAV, TAP and NA. Formal analysis: PV, CV and FC. Investigation: PV, JIE, CV, QKL, CTH, and FC, Resources: CIH, RFN, CTH, GF, DL and FC. Data Curation: PV and CAV. Writing–original draft: PV and FC. Writing – review & editing: PV, CV, RN, QKL, CTH, GF, DL, and FC. Funding acquisition: CIH, RFN, CTH, GF, DL and FC.

## Supplementary Information

Figure S1. Map of the world indicating where the *S. eubayanus* isolates were found. Each dot represents an isolation location. Distribution across Patagonia and the rest of the world is shown. Color code corresponds to different genetic lineages.

Figure S2. Samples and Isolation of *S. eubayanus* around the world. Eight countries from which *S. eubayanus* has been isolated. Pie charts above each bar represent the substrate from which samples were collected. Different colors in bar charts represent each population identified in this work.

Figure S3. Maximum likelihood tree of 471 *S. eubayanus* strains.

Figure S4. Cross-validation error value for the best K in fastStructure. The black dot represents the best value.

Figure S5. Plot of the genomic variation distribution of (A) 471 *S. eubayanus* strains based on the first two components of a PCA performed using 13,958 SNPs. (B) PCA considers only pure lineages, using 1,133,154 SNPs. Each dot represents a single strain.

Figure S6. Tajima’s *D* test statistic. Color codes in (B) and (C) represent each population identified in this work.

Figure S7. Model optimization for migration edges using TreeMix. TreeMix likelihood scores and percentage of variance explained across models with different numbers of migration edges (m). Each panel corresponds to an analysis with a different outgroup configuration: (A–B) *S. uvarum* as the outgroup, and (C) the PA population as the outgroup. Curves show the mean log-likelihood (± SD), variance explained, and the Δm statistic. Dashed lines indicate the 99.8% variance threshold used to select the optimal model

Figure S8. Structural Variation pairwise comparison. (A) SVs pairwise comparisons among all *S. eubayanus* genome assemblies with the total number of structural variations. (B) The range of total structural variation counts observed for each genome serves as a reference. Gray dots indicate each pairwise genome comparison. Colored dots indicate the mean and are colored by lineage. (C) The range of structural variation counts found for each type of variation. Gray dots indicate each pairwise genome comparison. Colored dots indicate the mean and are colored according to each lineage

Figure S9. Synteny of *MAL* genes across all *S. eubayanus* lineages. Genomic organization of the MAL loci on chromosome V (A) and chromosome XVI (B) across the reference strain CBS12357^T^, pure lineages (PA, PB1–PB5), and Hol-ADMIX strains (yHRVM108, UCD646, UCD650). Conserved synteny blocks are shown in grey.

Figure S10. Complementation of maltose growth deficiency in Hol-ADMIX strains. Growth performance of Hol-ADMIX strains UCD646 and yHRVM108, and their *MAL33*-complemented derivatives (UCD646_MAL33 and yHRVM108_MAL33), in media containing increasing maltose concentrations (5%, 10%, and 20%). Asterisks indicate statistically significant differences between wild-type and complemented strains (*p* < 0.05, *p* < 0.01, *p* < 0.001, *p* < 0.0001)

Figure S11. MAL33 complementation assay control. Growth of the control strain CL467 and its *MAL33*-complemented derivative (CL467_MAL33) in media containing 2% glucose or 2% maltose. Bars represent the maximum optical density (OD₆ ₀ ₀ ). Asterisks indicate statistically significant differences between strains under each condition (*p* < 0.01, *p* < 0.001)

**Supplementary Table 1.** List of *S. eubayanus* strains used in this study.

**Supplementary Table 2.** Varian Calling Summary statistics.

**Supplementary Table 3.** Population genetics summary statistics for each lineage.

**Supplementary Table 4.** Metrics for genome assemblies.

**Supplementary Table 5.** SVs pairwise comparison among all *S. eubayanus* genome assemblies.

**Supplementary Table 6.** SVs counts pairwise comparison among populations.

**Supplementary Table 7.** Phenotypic variation in maltose utilization among *S. eubayanus*.

**Supplementary Table 8.** Phenotypic variation in maltose utilization in complemented *S. eubayanus* strains with CBS12357 *MAL33* allele

**Supplementary Table 9.** Maltose concentration [µg/mg] in *Quercus* and *Nothofagus* trees determined in the dry bark samples.

